# Ptbp1 knockdown in mouse striatum did not induce astrocyte-to-neuron conversion using HA-tagged labeling system

**DOI:** 10.1101/2022.03.29.486202

**Authors:** Guixiang Yang, Zixiang Yan, Xiaoqing Wu, Meng Zhang, Chunlong Xu, Linyu Shi, Hui Yang, Kailun Fang

## Abstract

Conversion of astrocytes to neurons (AtN) is a promising potential strategy for the treatment of neurodegenerative diseases. Recent studies have reported that shRNA-, CasRx-, or ASO-mediated Ptbp1 suppression could reprogram resident astrocytes to neurons^1–3^. However, some groups have disputed the data interpretation of the reported AtN conversion events^4–7^. These controversies surrounding AtN conversion may due to differences in the astrocyte fate-mapping systems they applied from that in the original study, *i*.*e*., recombinant mouse strains with astrocyte specific reporter constructs versus AAV-based labeling systems. Here, we applied AAV-based tracing systems to label astrocytes with GFAP-driven HA-tagged Cas13X (AAV-GFAP::Cas13X-NLS-HA-sgPtbp1). Compared to GFAP-driven tdTomato reporter (AAV-GFAP::tdTomato) system in previous studies, we found no AtN conversion in mouse striatum. Furthermore, no intermediate neurons and no neuron density increase suggested no AtN conversion. Our findings indicated that inconsistent AtN outcomes may arise from different fate-mapping systems between AAV and transgenic mice, as well as through use of different reporter proteins. Thus, the complexity of astrocyte labeling systems warrants careful attention when drawing conclusions about whether AtN conversion occurs.

## Results

Previously, we found that CasRx could repress Ptbp1 in astrocytes and induce AtN conversion^2^. However, technical difficulties in the CasRx-mediated Ptbp1 knockdown reported by other groups^6^ led us to investigate alternative approaches to repress Ptbp1. To this end, we tested the efficiency of Ptbp1 knockdown by Cas13X, a hyper-compact CRISPR-Cas13 protein recently identified by our group^8^. We screened five sgRNAs targeting Ptbp1 mRNA (Figure S1A) and found that sgRNAs-2, -3 and -5, independently or combined, could effectively knock down Ptbp1 expression in HEK293T, Cos7 and N2a cell lines (Figure S1B).

To knockdown Ptbp1*in vivo*, we applied an AAV-PHP.eB capsid with a 681 bp-length human GFAP promoter (hGFAP)^9^ to drive astrocyte-specific expression of Cas13X-NLS-HA-sgPtbp1-(2,3,5) or the non-target control (sgNT) in C57BL/6 mice (Figure 1A). Immunofluorescent staining of brain sections showed that the PTBP1 signal significantly decreased over time (Figure 1B, S1C, and S1D), indicated by the decreasing proportion of Ptbp1+ HA+ GFAP+ astrocytes in AAV-hGFAP::Cas13X-sgPtbp1 treated mice from 74.79% at 1 week post-injection to 18.42%, 14.96%, 11.98% at 2 weeks, 1 month, and 2 months post-injection, respectively (Figure 1C). By contrast, the proportion of Ptbp1+ HA+ GFAP+ astrocytes in AAV-hGFAP::Cas13X-sgNT treated mice was not significantly changed from 1 week to 2 months post-injection (Figure 1B and 1C).

**Figure 1.**
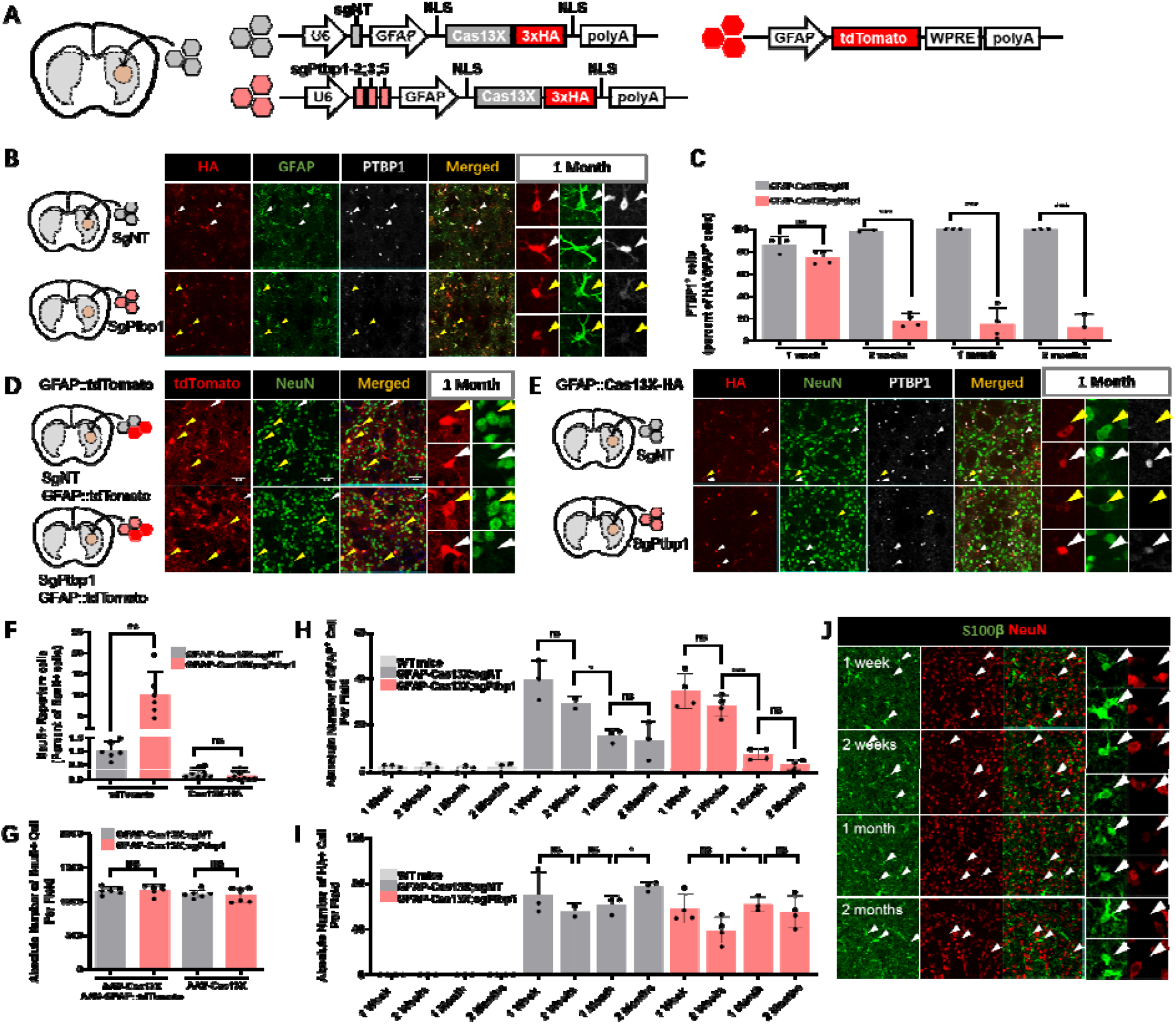
Use of different tracing markers leads to discrepancies in AtN conversion data in mouse striatum under Ptbp1 knockdown discrepancies in astrocyte labeling. **(A)** Schematic of AAV-GFAP::Cas13X-sgNT, AAV-Cas13X-sgPtbp1-2;3;5, and AAV-GFAP::tdTomato constructs. NT: non-targeted. **(B)** Representative images of striatum sections at 1 month after injection (1.5E9 vg/striatum) with immunofluorescent staining for HA, GFAP, and Ptbp1. White arrowheads: cells without Ptbp1 suppression. Yellow arrowheads: Ptbp1-suppressed cells. **(C)** Quantification and statistical analysis of Ptbp1+ cell proportions among HA+GFAP+ cells, indicating Ptbp1 knockdown efficiency in AAV-transduced astrocytes. **(D)** Representative images of brain sections with immunofluorescent staining for NeuN at 1 month after injection. Yellow arrowheads: tdTomato+ NeuN+ cells. White arrowheads: tdTomato+ NeuN-cells. **(E)** Representative images of brain sections stained for HA, NeuN, and Ptbp1 at 1 month post-injection. Yellow arrowheads: HA+ NeuN+ cells. White arrowheads: HA+ NeuN-cells. **(F)** Quantification and statistical analysis of tdTomato+ NeuN+ or HA+ NeuN+ cell proportions among NeuN+ cells, at 1 and 2 months post-injection. **(G)** Absolute numbers of NeuN+ cells at 1 and 2 months after injection with AAV-Cas13X or AAV-Cas13X plus AAV-GFP::tdTomado. **(H)** Absolute number of endogenous GFAP+ cells at 1, 2, 4, and 8 weeks post-injection. **(I)** Absolute numbers of exogenous HA+ cells at 1, 2, 4, and 8 weeks post-injection. **(J)** Representative images of brain sections at different time points. White arrowheads: S100β+ NeuN-cells. WT mice, wild-type littermate mice without AAV treatment. Significance was determined by two-tailed t test. * P<0.05, ** P<0.001, *** P<0.0001, ns: not significant.

To determine whether Ptbp1 suppression could successfully induce AtN conversion, the Ptbp1-knockdown virus (AAV-hGFAP::Cas13X-sgPtbp1) and astrocyte-tracing virus (AAV-hGFAP::tdTomato) were co-injected into the striatum of wild-type (WT) mice (Figure 1D). At 1 month (Figure 1D) and 2 months (Figure S1E, 1F) after injection, the populations of tdTomato+ NeuN+ cells were significantly greater in the Ptbp1 knockdown group compared to that in non-target control group (10.04% vs 1.02% tdTomato+ among total NeuN+ cells, respectively; Figure 1F). These results indicated potential AtN conversion, which was consistent with observations in several previous studies^2,3^. However, the NeuN+ cell density was not significantly changed, which indicated that further validation was needed to determine whether AtN conversion occurred (Figure 1G).

To further confirm the AtN conversion observed by tdTomato+/NeuN+ labeling, we then used an HA tag in hGFAP::Cas13X-NLS-HA-sgPtbp1 construct to track astrocytes at 1 month and 2 months post-injection. Immunofluorescent staining revealed that HA+ NeuN+ cell populations were not significantly different between Ptbp1 knockdown and control mice (0.14% in Ptbp1 KD group vs 0.22% in non-target controls, P=0.2157, Figure 1E, 1F, S1G, and S1H). Since AtN intermediate cells might emerge earlier than 1 month, we also tracked the HA signal at 1 week and 2 weeks post-injection, in the early stages of Ptbp1 suppression. Again, no significant differences in HA+ NeuN+ cell proportions could be detected between Ptbp1 knockdown and non-target control group (Figure S1G, and S1H), which aligned well with several recent reports of no AtN conversion in transgenic mouse models^4–7^. In addition, NeuN+ cell density was not significantly changed (Figure 1G). Meanwhile, not like NeuroD1 induced AtN conversion^10^, we observed no S100β+ NeuN+ AtN intermediate cells among thousands of S100β+ Ptbp1^low^ astrocytes from 1 week to 2 months post-injection (Figure 1J), suggesting that AtN conversion did not occur.

## Discussion

It is controversial as to whether astrocytes can be converted to neurons after Ptbp1 knockdown. Here we observed no significant changes in the proportion of HA+NeuN+ cells, no changes in NeuN+ cell density, and no intermediate conversion cells in the Ptbp1 knockout group when using HA-tagged labeling system, indicating that no AtN conversion occurred. Although significant more TdTomato+ NeuN+ cells have been found in the Ptbp1-suppressed striatum, this result may be misled by the related astrocyte-tracing systems (AAV-pGFAP::TdTomato).

In this previous studies and this work, different astrocyte-tracing systems showed the opposite AtN results. One difference from these astrocyte-labeling systems is that different astrocyte-specific promoters were applied. The GFAP promoter is the common promoter for astrocyte labeling. However, the introduction of GFAP promoter is well-known to result in high levels of leaky expression in neurons, up to 50%^6^. Meanwhile, aberrant activity by the GFAP promoter has been observed under certain conditions ^6,11,12^, suggesting GFAP promoter is not appropriate for astrocyte-labeling. To restrict the astrocyte labeling specificity, the Aldh1l1 promoter is currently applied. Since the Aldh1l1-CreER^T2^;LSL-YFP transgenic mouse lines exhibit high specificity in astrocyte, with relatively low (4.3%) neuron mislabeling, it is more specific than GFAP promoter^6,13^. Indeed, three independent groups using Aldh1l1-CreER^T2^ labeling systems and observed no AtN conversion through Ptbp1 knockdown or knockout *in vivo* ^4,6,7^. However, application of an endogenous promoter-driven mGFAP-Cre;LSL-YFP mouse strain enabled stringent, astrocyte-specific YFP expression, which also showed no AtN conversion under Ptbp1 knockdown^6^. Together, these findings indicated that GFAP promoter alone cannot fully account for the non-specific reporter signals in astrocyte labeling experiments.

Another explanation account for the conflicting results could be differences between exogenous human GFAP promoter (delivered by AAV vector) and endogenous mouse GFAP promoter (in mouse strain). The endogenous GFAP promoter in mouse genome is longer and harbors different recognition sites from that of the truncated exogenous hGFAP promoter delivered by AAV, and is thus subject to control by different regulatory elements^14–17^. In our studies, the endogenous GFAP-driven signal continuously decreases from 1 week to 2 months in mice *in vivo* (Figure 1H), whereas the HA signal driven by an exogenous hGFAP promoter persisted for 2 months (Figure 1I), indicating distinct differences between the mouse and AAV-mediated GFAP promoter activity. Moreover, under AAV infection of neurons, several copies of the AAV genome harboring exogenous GFAP promoters enter a single cell, which could result in different patterns of expression from that driven by one or two copies of endogenous GFAP promoter in mouse. This phenomenon could account for the significantly greater labeling of reporter+ neurons using AAV than using mouse strain^6^. Meanwhile, when removing WPRE element of AAV-GFAP::tdTomato labeling system to low down tdTomato signal, no AtN conversion could be observed in a more stringent astrocyte labeling system in retina in a recent study^18^.

In addition, in this work we also demonstrated that different reporters, such as our tests of an AAV-based hGFAP-tdTomato and an –HA tag, can generate markedly different outcomes in AtN conversion labeling under Cas13X-mediated Ptbp1 knockdown. These disparities suggest that we cannot exclude the effects of different reporter proteins in addition to the leakiness of the GFAP promoter in conflicting astrocyte labeling experiments. Previous reports have shown that endogenously-driven CRE could be transferred into neighboring neurons and there induce reporter expression through exosomes or tunneling nanotubes^19,20^. In the current study, it is possible that some excess hGFAP::tdTomato protein could be transferred from astrocytes to adjacent neurons in a manner similar to CRE through an as-of-yet undetermined route. This explanation could account for the observed leakage of tdTomato signal in neurons, in addition to promoter driven leaky expression. It would align well with observations by two conflicting studies, one reported high astrocyte specificity of an GFAP-CreER^T2^;LSL-YFP mouse strain tracing system with no detectable AtN conversion^6^, but in another study, AtN was observed using GFAP-CreERTM;Rosa-Tdtomato mouse strain^3^. One of the differences between the two studies is the reporter protein.

In addition, there are some reports using transgene expression or Ptbp1 knockdown treatment showing significant improvements in animal models of neurodegenerative disease. These may have explanations other than AtN conversion, including rejuvenation of endogenous neurons. For example, overexpression of Nmnat1 could protect retinal ganglion cells in mouse models of glaucoma^21^. For these phenotypic changes in AtN studies, other explanations should be considered.

Therefore, our findings imply no AtN coversion in Ptbp1 suppressed mouse striatum. Data showing *in vivo* neuronal lineage conversion obtained by reporter proteins such as tdTomato should be taken with caution and subjected to validation by additional reporters (*e*.*g*., HA tag, YFP, SUN1-GFP etc.) regardless of whether a viral vector or transgenic mouse line was used to label the fate-mapping.

## SUPPLEMENTAL FIGURE LEGEND

**Figure S1.**
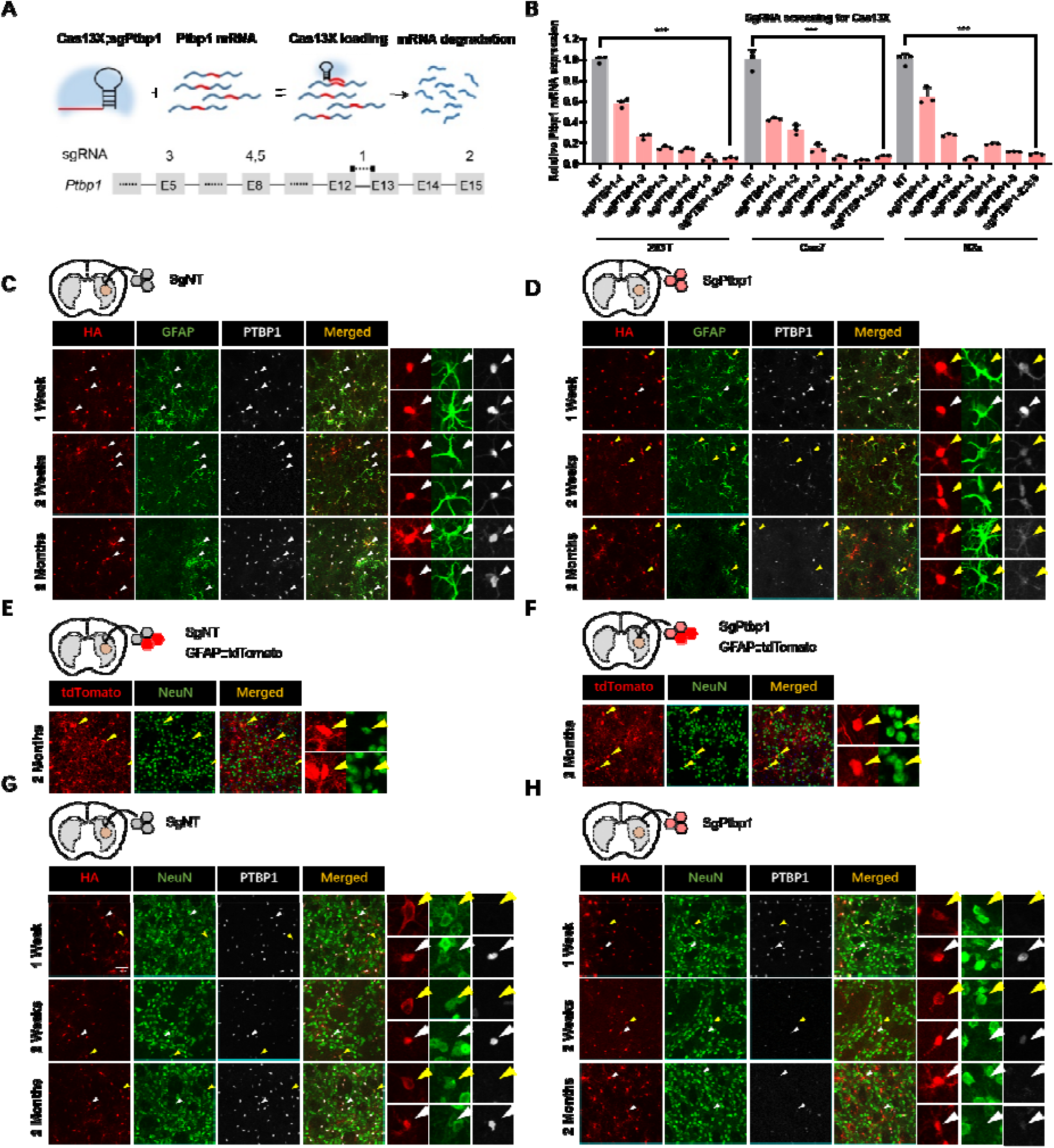
Additional time points showing differences in astrocyte labeling specificity and astrocyte-to-neuron conversion data between tdTomato and HA tag reporters. **(A)** Schematic of Cas13X mode of knockdown and Ptbp1 sgRNA design targeting exons 5, 8, 12, 13 and 15; **(B)** Relative mRNA expression of Ptbp1 under treatments with Cas13X;sgPtbp1 or Cas13X;sgNT in 293T, Cos7, and N2a cell lines. ***: P value<0.0001 in two-tailed t-test.**(C**,**D)** Representative images of brain sections at 1, 2, and 8 weeks after non-target control AAV or Ptbp1 knockdown virus injection. Stained for HA, GFAP, and PTBP1.**(E**,**F)** Representative images of brain sections at 2 months after non-target control AAV or Ptbp1 knockdown virus injection, co-injected with AAV-GFAP::tdTomato to label astrocyte.**(G**,**H)** Representative images of brain sections at 1, 2, and 8 weeks after non-target control AAV or Ptbp1 knockdown virus injection. Stained for HA, NeuN, and PTBP1.

## MATERIALS AND METHODS

### Vector and gRNA sequences

The vector information is provided in Supplementary Sequences. SgPtbp1-2:

5’-tgtggttggagaactggatgtagatgggct-3’; sgPtbp1-3: 5’-gagcccatctggatcagtgccatcttgcgg-3’; sgPtbp1-5: 5’-agtcgatgcgcagcgtgcagcaggcgttgt-3’; sgNT: 5’-attggcaccatgccgtgggtttcaatattg-3’.

### Cell culture and plasmids transfection

N2a, Cos7 and 293T cell lines were obtained from Cell bank of Shanghai Institute of Biochemistry and Cell Biology (SIBCB), Chinese Academy of Sciences (CAS). They were cultured in 37□ and 5% CO2 incubator in medium DMEM (Gibco) supplemented with 10% fetal bovine serum (FBS, Gibco) and 1 × penicillin / streptomycin (Gibco). 3 μg plasmids were transiently transfected with 6 μl Polyethylenimine (PEI) into cultured cells in 12-well clusters.

### RNA extraction and RT-qPCR

The transfected plasmids containing co-expression elements of mCherry. Three days after transfection, mCherry-positive cells (top 20%) were collected by fluorescence activated cell sorting (FACS). RNA was extracted with TRIzol Reagent (Ambion). cDNA were obtained with HiScript Q RT SuperMix for qPCR Kit (Vazyme). Quantitative PCR (qPCR) was performed using AceQ qPCR SYBR Green Master Mix (Vazyme) and LightCycler 480 II (Roche).

### Mice

10-week-old male C57BL/6 mice were obtained from Vital River Laboratory Animal Technology Co., Ltd. All animal experiments were performed and approved by the Animal Care and Use Committee of the Institute of Huigene Therapeutics Inc., Shanghai, China.

### AAV package and preparation

Viral particles of AAV-PHP.eB were packaged in co-transfected HEK293T cells with the other two plasmids: pAAV-Rep-Cap and pAAV-Helper. After harvest, viral particles were purified with a heparin column (GE HEALTHCARE BIOSCIENCES) and then concentrated with an Ultra-4 centrifugal filter unit (Amicon, 100,000 molecular weight cutoff). Titers of viral particles were determined by qPCR to achieve >1E12 particles/ml.

### Striatum injection

10-week-old male C57BL/6 mice were anaesthetized with Zoletil® 50 (Virbac, 0.1ml/10g) and then placed in a stereotaxic mouse frame. The skin over the skull was shaven and opened using a razor. 1.5μl of AAV was injected into the striatum at the following coordinates (relative to bregma): anteroposterior (A/P) = +0.75 mm, mediolateral (M/L) = −1.9 mm, dorsoventral (D/V) = −3.45 mm. The viral solution was injected slowly (300nl/min).

### Immunofluorescence

Immunofluorescence staining was performed at different time points after AAV injection. The brains were perfused and fixed with 4% paraformaldehyde (PFA) overnight and kept in 30% sucrose for at least 12 hours. Brains were sectioned after embedding and freezing, and slices with the thickness of 30 mm were used for immunofluorescence staining. Brain sections were rinsed thoroughly with 0.1 M phosphate buffer (PB). Primary antibodies: Rabbit anti-PTBP1(1:500, PA5-81297, Invitrogen), Rabbit anti-NeuN(1:500, 24307S, Cell Signaling Technology), Mouse anti-NeuN(1:500, AB104224, Abcam), Rat anti-HA(1:500, 11867423001, ROCHE), Mouse anti-GFAP(1:500, AB279290, Abcam), Mouse anti-S100β(1:500, S2532, Merck). Secondary antibodies: Alexa Fluor® 488 AffiniPure Donkey Anti-Rabbit IgG (H+L) (1:1000, 711-545-152, Jackson ImmunoResearch Labs), Alexa Fluor® 488 AffiniPure Donkey Anti-Mouse IgG (H+L)(1:1000, 715-545-151, Jackson ImmunoResearch Labs), Cy™5 AffiniPure Donkey Anti-Rat IgG (H+L)(1:1000, 712-175-153, Jackson ImmunoResearch Labs), Cy3 AffiniPure Donkey Anti-Rabbit IgG (H+L)(1:1000, 711-165-152, Jackson ImmunoResearch Labs) were used in this study. After antibody incubation, slices were washed and covered with mountant (Life Technology). Images were visualized under a NIKON C2+ microscope.

## ACKNOWLEDGMENTS

We thank Bai Weiya and Geng Guannan for AAV vector preparation, Gao Yanxia and Cheng Leping for project discussion. This work was supported by Chinese National Science and Technology major project R&D Program of China (2017YFC1001302 and 2018YFC2000101), Strategic Priority Research Program of Chinese Academy of Science (XDB32060000), National Natural Science Foundation of China (31871502, 31901047, and 82021001), Basic Frontier Scientific Research Program of Chinese Academy of Sciences From 0 to 1 original innovation project (ZDBS-LY-SM001), Lingang Lab (Grant LG202106-01-02), and International Partnership Program of Chinese Academy of Sciences (153D31KYSB20170059).

## Antibodies

| **Antibodies** | **SOURCE** | **Cat.** |
| --- | --- | --- |
| Rabbit anti-PTBP1 | Invitrogen | PA5-81297 |
| Rabbit anti-NeuN | Cell Signaling Technology | 24307S |
| Mouse anti-NeuN | Abcam | AB104224 |
| Rat anti-HA | ROCHE | 11867423001 |
| Mouse anti-GFAP | Abcam | AB279290 |
| Mouse anti-S100β | Merck | S2532 |
| Alexa Fluor® 488 AffiniPure Donkey Anti-Rabbit IgG (H+L) | Jackson ImmunoResearch Labs | 711-545-152 |
| Alexa Fluor® 488 AffiniPure Donkey Anti-Mouse IgG (H+L) | Jackson ImmunoResearch Labs | 715-545-151 |
| Cy<sup>TM</sup>5 AffiniPure Donkey Anti-Rat IgG (H+L) | Jackson ImmunoResearch Labs | 712-175-153 |
| Cy3 AffiniPure Donkey Anti-Rabbit IgG (H+L) | Jackson ImmunoResearch Labs | 711-165-152 |

## AAV Titer and Dose

| Vectors and Virus | Titer | Work Titer | Volume (μl) |
| --- | --- | --- | --- |
| AAV-PHP.eB-hGFAP::Cas13X-NLS-HA-sgPtbp1-(2,3,5) | 5.47E+13 vg/ml | 1.0E+12 vg/ml | 1.5 |
| AAV-PHP.eB-GFAP-Tdtomato | 5.30E+13 vg/ml | 6.7E+11 vg/ml |  |
| AAV-PHP.eB-hGFAP::Cas13X-NLS-HA-sgNT | 1.23E+14 vg/ml | 1.0E+12 vg/ml | 1.5 |
| AAV-PHP.eB-GFAP-Tdtomato | 5.30E+13 vg/ml | 6.7E+11 vg/ml |  |
| AAV-PHP.eB-hGFAP::Cas13X-NLS-HA-sgPtbp1-(2,3,5) | 5.47E+13 vg/ml | 5.0E+12 vg/ml | 1.5 |
| AAV-PHP.eB-hGFAP::Cas13X-NLS-HA-sgPtbp1-(2,3,5) | 5.47E+13 vg/ml | 1.0E+12 vg/ml | 1.5 |
| AAV-PHP.eB-hGFAP::Cas13X-NLS-HA-sgNT | 1.23E+14 vg/ml | 5.0E+12 vg/ml | 1.5 |
| AAV-PHP.eB-hGFAP::Cas13X-NLS-HA-sgNT | 1.23E+14 vg/ml | 1.0E+12 vg/ml | 1.5 |

## AUTHOR CONTRIBUTIONS

L.S., H.Y., and K.F. designed the research, G.Y., Z.Y., X.W., M.Z. performed experiments and analyzed data. C.X., H.Y. and K.F. wrote the manuscript with input from all authors.

## REFERENCES

1. Maimon, R., Chillon-Marinas, C., Snethlage, C.E., Singhal, S.M., McAlonis-Downes, M., Ling, K., Rigo, F., Bennett, C.F., Da Cruz, S., Hnasko, T.S., Muotri, A.R., et al. (2021). Therapeutically viable generation of neurons with antisense oligonucleotide suppression of PTB. Nature neuroscience 24, 1089–1099. 10.1038/s41593-021-00864-y.

2. Zhou, H., Su, J., Hu, X., Zhou, C., Li, H., Chen, Z., Xiao, Q., Wang, B., Wu, W., Sun, Y., Zhou, Y., et al. (2020). Glia-to-Neuron Conversion by CRISPR-CasRx Alleviates Symptoms of Neurological Disease in Mice. Cell. 10.1016/j.cell.2020.03.024.

3. Qian, H., Kang, X., Hu, J., Zhang, D., Liang, Z., Meng, F., Zhang, X., Xue, Y., Maimon, R., Dowdy, S.F., Devaraj, N.K., et al. (2020). Reversing a model of Parkinson’s disease with in situ converted nigral neurons. Nature 582, 550–556. 10.1038/s41586-020-2388-4.

4. Hoang, T., Kim, D.W., Appel, H., Pannullo, N.A., Leavey, P., Ozawa, M., Zheng, S., Yu, M., Peachey, N.S., Kim, J., and Blackshaw, S. (2021). Ptbp1 deletion does not induce glia-to-neuron conversion in adult mouse retina and brain. 2021.2010.2004.462784. 10.1101/2021.10.04.462784 %J bioRxiv.

5. Leib, D., Chen, Y.H., Monteys, A.M., and Davidson, B.L. (2022). Limited astrocyte-to-neuron conversion in the mouse brain using NeuroD1 overexpression. Molecular therapy : the journal of the American Society of Gene Therapy 30, 982–986. 10.1016/j.ymthe.2022.01.028.

6. Wang, L.L., Serrano, C., Zhong, X., Ma, S., Zou, Y., and Zhang, C.L. (2021). Revisiting astrocyte to neuron conversion with lineage tracing in vivo. Cell. 10.1016/j.cell.2021.09.005.

7. Chen, W., Zheng, Q., Huang, Q., Ma, S., and Li, M. (2021). Repressing PTBP1 is incapable to convert reactive astrocytes to dopaminergic neurons in a mouse model of Parkinson’s disease. 2021.2011.2012.468309. 10.1101/2021.11.12.468309 %J bioRxiv.

8. Xu, C., Zhou, Y., Xiao, Q., He, B., Geng, G., Wang, Z., Cao, B., Dong, X., Bai, W., Wang, Y., Wang, X., et al. (2021). Programmable RNA editing with compact CRISPR-Cas13 systems from uncultivated microbes. Nature methods 18, 499–506. 10.1038/s41592-021-01124-4.

9. Lee, Y., Messing, A., Su, M., and Brenner, M. (2008). GFAP promoter elements required for region-specific and astrocyte-specific expression. Glia 56, 481–493. 10.1002/glia.20622.

10. Zheng, J., Li, T., Qi, S., Qin, B., Yu, J., and Chen, G. (2022). Neuroregenerative gene therapy to treat temporal lobe epilepsy in a rat model. Prog Neurobiol 208, 102198. 10.1016/j.pneurobio.2021.102198.

11. Su, M., Hu, H., Lee, Y., d’Azzo, A., Messing, A., and Brenner, M. (2004). Expression specificity of GFAP transgenes. Neurochem Res 29, 2075–2093. 10.1007/s11064-004-6881-1.

12. Hol, E.M., Roelofs, R.F., Moraal, E., Sonnemans, M.A., Sluijs, J.A., Proper, E.A., de Graan, P.N., Fischer, D.F., and van Leeuwen, F.W. (2003). Neuronal expression of GFAP in patients with Alzheimer pathology and identification of novel GFAP splice forms. Mol Psychiatry 8, 786–796. 10.1038/sj.mp.4001379.

13. Srinivasan, R., Lu, T.Y., Chai, H., Xu, J., Huang, B.S., Golshani, P., Coppola, G., and Khakh, B.S. (2016). New Transgenic Mouse Lines for Selectively Targeting Astrocytes and Studying Calcium Signals in Astrocyte Processes In Situ and In Vivo. Neuron 92, 1181–1195. 10.1016/j.neuron.2016.11.030.

14. Miura, M., Tamura, T., and Mikoshiba, K. (1990). Cell-specific expression of the mouse glial fibrillary acidic protein gene: identification of the cis-and trans-acting promoter elements for astrocyte-specific expression. J Neurochem 55, 1180–1188. 10.1111/j.1471-4159.1990.tb03123.x.

15. Sarkar, S., and Cowan, N.J. (1991). Intragenic sequences affect the expression of the gene encoding glial fibrillary acidic protein. J Neurochem 57, 675–684. 10.1111/j.1471-4159.1991.tb03799.x.

16. Besnard, F., Brenner, M., Nakatani, Y., Chao, R., Purohit, H.J., and Freese, E. (1991). Multiple interacting sites regulate astrocyte-specific transcription of the human gene for glial fibrillary acidic protein. The Journal of biological chemistry 266, 18877–18883.

17. Nakatani, Y., Horikoshi, M., Brenner, M., Yamamoto, T., Besnard, F., Roeder, R.G., and Freese, E. (1990). A downstream initiation element required for efficient TATA box binding and in vitro function of TFIID. Nature 348, 86–88. 10.1038/348086a0.

18. Gao, Y., Fang, K., Yan, Z., Zhang, H., Geng, G., Wu, W., Xu, D., Zhang, H., Zhong, N., Wang, Q., Cai, M., et al. (2022). Develop an efficient and specific AAV based labeling system for Muller glia in mice. 2021.2012.2010.472182. 10.1101/2021.12.10.472182 %J bioRxiv.

19. Fruhbeis, C., Frohlich, D., Kuo, W.P., and Kramer-Albers, E.M. (2013). Extracellular vesicles as mediators of neuron-glia communication. Front Cell Neurosci 7, 182. 10.3389/fncel.2013.00182.

20. Wang, Y., Cui, J., Sun, X., and Zhang, Y. (2011). Tunneling-nanotube development in astrocytes depends on p53 activation. Cell Death Differ 18, 732–742. 10.1038/cdd.2010.147.

21. Williams, P.A., Harder, J.M., and John, S.W.M. (2017). Glaucoma as a Metabolic Optic Neuropathy: Making the Case for Nicotinamide Treatment in Glaucoma. J Glaucoma 26, 1161–1168. 10.1097/IJG.0000000000000767.

